# Redistribution of proteasomes through diffusion and cytoskeleton-dependent mechanisms upon stress induced by protein aggregates

**DOI:** 10.1101/487702

**Authors:** Michael J Morten, Yu Zhang, Bing Li, Jonathan X Meng, Kun Jiang, Liina Sirvio, Ji Eun Lee, Anna Helena Lippert, Matilda Burridge, Katiuska Daniela Pulgar Prieto, Alexander Roy Carr, Aleks Ponjavic, Steven F Lee, Daniel Finley, David Klenerman, Yu Ye

## Abstract

Proteasomes are abundant molecular machines distributed throughout the eukaryotic cell to facilitate protein degradation. We previously showed that proteasomes can re-organize and assemble into foci bodies in response to proteotoxic stress^1^, which we termed transient aggregate-associated droplets (TAADs)^2^. Here, we use single-molecule localization microscopy to quantify re-organization of mammalian proteasomes in different subcellular regions during proteotoxic stress to engage with invading protein aggregates. We determine that ∼60% of 20S proteasomes are capped by at least one 19S particle, and this ratio remains constant during stress. Using single-particle tracking, we show that cells confine global proteasomal movement when experiencing proteotoxic stress via a cytoskeleton-dependent mechanism. Similar limitations on proteasome movement are also observed upon membrane depolarization and repolarization events by patch clamp, which directly induce proteasomes to move away or towards the plasma membrane, respectively. Together, our experiments reveal distinct modes of proteasome motion dependent on cellular requirements, and propose that proteasome transport is restricted upon aggregate invasion to facilitate their subsequent degradation.

## Introduction

The ubiquitin-proteasome system controls the abundance of cellular proteins by selective degradation and thereby regulating key signaling processes, such as oxidative stress, inflammatory and unfolded protein responses^3–5^. Ubiquitin-dependent degradation is canonically carried out by the 26S proteasome holoenzyme, which is assembled from 20S core-(CP) and 19S regulatory particles (RP). The holoenzyme is believed to be the preferred proteasome assembly, and may reversibly disassemble into CPs and RPs upon altered cellular requirements^6^ (**SFig.1A**). Free CPs are thought to degrade unfolded and disordered proteins, including nascent protein turnover during neuronal stimulation^7^, while free RPs have recently been found with increased abundance at the synapse and suggested to regulate synaptic transmission^8^. Another recent report provided evidence for ubiquitin-independent selective degradation of particularly nuclear proteins and transcription factors by the holoenzyme^9^, further underscoring the versatile functions and regulatory roles assigned to proteasome particles.

Earlier studies have reported that proteasomes are present throughout the cell^10–12^, presumably to facilitate local protein turnover, though the observed relative distribution can vary between biological conditions and imaging methods^13–15^. Altered physiological conditions such as hypertonic stress, osmotic shock and starvation have been shown to induce re-distribution of proteasomes into foci bodies, which have been given multiple names, including clastosomes, senescence-associated nuclear proteasome foci (SANPs), and proteasome storage granules (PSGs)^16–20^. Such foci are thought to help rebalance proteostasis in response to various types of cell stress; PSGs are associated with a protective role to sequester proteasomes during starvation^21^, and clastosomes may function as degradation centers in response to hypertonic stress^19^. More recently, we postulated that certain foci types may serve to concentrate proteasomal co-proteins and enzymes of the ubiquitin system to facilitate disaggregation and degradation, which we termed transient aggregate-associated droplets (TAADs)^2^ and provided evidence for their induction by protein aggregates^1^. The formation of such TAADs requires re-organization of cellular proteasomes, though the cellular mechanisms enabling their formation are not well-studied.

In this work, we used single-molecule localization microscopy (SMLM) to study the redistribution of proteasomes in response to toxic protein aggregates that have entered the cell. Quantifying the subcellular distribution and the ratio of the different proteasome particles did not detect any significant stress-induced changes. We further showed that both TAAD formation and proteasome movement induced by membrane de-/hyperpolarization also involve cytoskeleton-dependent mechanisms. Critically, TAAD formation was accompanied by a global reduction of freely diffusing proteasomes and an increase in the confinement of proteasomes throughout the cytoplasm, as observed by single-particle tracking (SPT). Together, our data suggest that proteasomes are dynamic particles that change their mode of motion, cellular distribution and local concentration in response to external stimuli as required by the cell.

## Results

### Proteotoxic and hypertonic stress induce redistribution of proteasomes

To study proteasome dynamics in cells, we edited the genomic sequences of PSMD14/RPN11 or PSMB2/PRE1, subunits of the RP or CP, respectively, to be expressed in tandem with fluorescent protein (FP) sequences encoding eGFP and mEos (**Fig.1A** and **SFig.1A-B**, see **Materials and Methods**). The CRISPR-edited monoclonal cell lines were confirmed by genotyping and incorporation of FP-tagged subunits into proteasomal particles was verified by native gels (**SFig.1C-D**), in line with previous studies^17,22,23^. The ratios of FP-tagged to untagged proteasomes were determined by densitometry analysis from Western blots (**SFig.2**).

**Figure 1.**
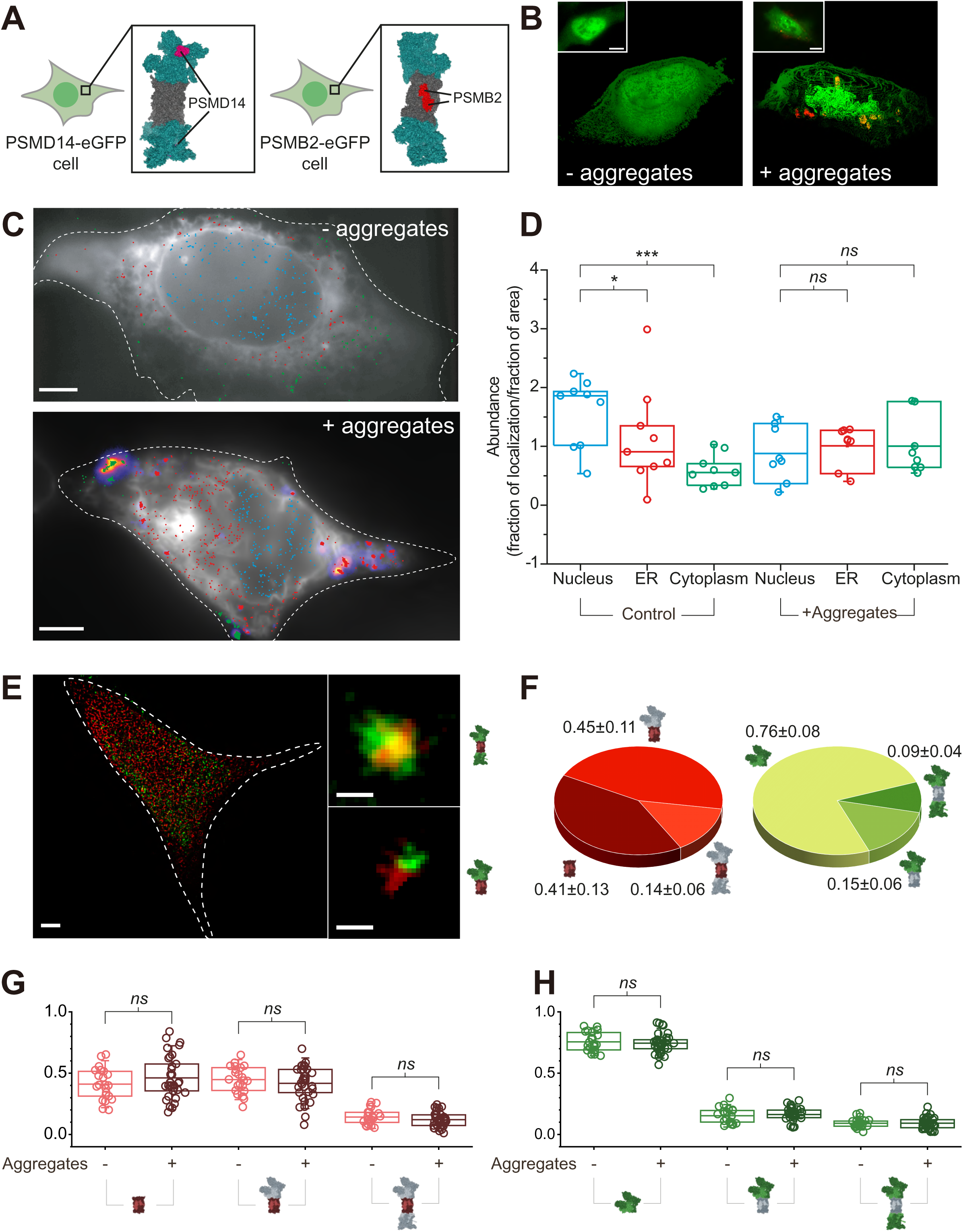
Proteotoxic stress induces major re-distribution of proteasome particles. (**A**) Cartoon representation of CRISPR-edited cells expressing subunit of the RP (PSMD14, *left*) or CP (PSMB2, *right*) in tandem with FP at the C-terminus. (**B**) A typical PSMD14-eGFP cell reconstructed in 3D from HILO imaging with 100 nm step size z-stack before (*left*) and 24 hrs after (*right*) incubation with aS aggregates (Alexa647 partial labeling). Scale bar = 5 μm. (**C**) HILO imaging of individual proteasomal particles in PSMD14-mEos cells expressing an ER marker, sec61-mEos (white), at rest versus after 24 hrs incubation with aS-Alexa647 (fire-LUT). Cells were imaged using SMLM (see **Materials and Methods**). Reconstructed RPs localized to the nucleus (blue), overlapping with the ER (cyan), or cytosol (green) are color-coded. Scale bars = 1 µm. (**D**) Calculated relative abundance of PSMD14-mEos found in each subcellular compartment before and after aS aggregate incubation presented in Box-plots (n = 9 and n = 8 cells, respectively). Statistical significance was calculated using student’s t-test. (**E**) Immunofluorescence labeling and SMLM imaging of a typical HEK293A cell for CP (red) and RP (green) enabled calculation of capped versus uncapped proteasome ratios. Scale bars represent 5 μm and 100 nm inset. Inset with cartoon representations of singly- and doubly capped holoenzymes, are shown (*right*). (**F**) Relative ratios of free CP and RP versus singly- and doubly capped holoenzymes in HEK293A are shown in pie charts. Mean ± standard deviations from 22 cells with uncertainties are presented to one significant figure, and each mean value is reported to the same decimal place as its error. (**G-H**) HEK293A cells were incubated with aS aggregates for 24 hrs (n = 33 cells) and labeled as in **E,** for fractional quantification of (**G**) CP and (**H**) RP. Box-plots show that incubation of aggregates does not induce any significance changes between the relative ratios of free particles or holoenzymes. This is in line with observations made in differentiated SH-SY5Y neuroblastoma cells (**SFig.5**).

We examined the spatial distribution of proteasomes using both total-internal reflection fluorescence (TIRF)/HILO^24^ and light-sheet microscopy, confirming that both CP and RP were present throughout the cell, with strong presence in the nucleus (**Fig.1B** *left*, **SFig.3A** *left,* and **SFig.3B** *top*). Significant proteasomal re-distribution was observed upon proteotoxic stress, induced by the treatment of pre-assembled recombinant alpha-synuclein (aS) aggregates that were partially labeled with Alexa Fluor 647, or Alexa647 (**Fig.1B** *right*, **SFig.3A** *right* and **SFig.3B** *bottom*). In line with our previous report^1^, strong fluorescence intensities from proteasome particles were identified in close proximity to the internalized aggregates and consistent with our description proposed for TAADs^2^. To quantify proteasome re-distribution for the observed TAAD formation, we first imaged PSMD14- or PSMB2-mEos cells by SMLM and counted the number of proteasomes either found within the nucleus, co-localizing with the ER, or residing elsewhere in the cytosol (see **Materials and Methods**). In unstressed cells, proteasomes were densest in the nucleus and gradually reduced in density at the ER and the cytosol (**Fig.1C-D**). This density gradient was shifted by aggregate-induced proteotoxic stress, which caused significantly increased proteasome levels in the cytosol at the expense of reduced levels in the nucleus. We interpreted these two distributions as a resting and a stressed state, where cells responded to invading aggregates by reorganizing proteasomes across the subcellular compartments. For comparison, we assembled hypertonic-induced foci by adding CaCl_2_ to the cell media. These foci were mostly localized in the nucleus, and distinct to aggregate-induced proteasome relocalization (**SFig.4A-B**). This suggests that proteasome distribution is dynamically associated with the type of stress signal.

To examine if proteasome assemblies were altered by stress, we quantified the level of free CP and RP versus singly- or doubly-capped proteasome holoenzymes using SMLM at high-precision (∼4 nm)^1^ (**Fig.1E** and **Materials and Methods**). In resting cells, we observed high correlation between CPs and RPs, with the majority of CP associated with one (0.45±0.11) or two (0.14±0.06) RP and therefore forming singly- or doubly-capped holoenzymes, respectively (**Fig.1F**, *left*). A significant fraction (0.41±0.13) of CP was detected without association with RP and therefore classified as free particles. Repeating the relative quantification for RPs, we observed similar levels of RPs forming part of singly-(0.15±0.06) and doubly-(0.09±0.04) capped holoenzymes, while most RPs (0.76±0.07) remained as free particles (**Fig.1F**, *right*). The observed differences in relative free CP and free RP ratios are consistent with a recent observation in neurons^8^, and suggest that the majority of CP activities are coupled to those of the RP, while RPs themselves may have distinct functions, e.g. as deubiquitinases^8^ or chaperones of misfolded proteins^25^. Consistently, we observed similar distributions of CPs (0.32±0.18, 0.22±0.19, 0.46±0.15 for singly-, doubly-capped and free particles, respectively) and RPs (0.25±0.16, 0.31±0.26, 0.45±0.20) in differentiated SH-SY5Y neuroblastoma cells (**SFig.5A-B**). Intriguingly, proteotoxic stress by aS aggregates did not quantitatively change the relative ratios of free CPs and RPs in either cell types (**Fig.1G-H** and **SFig.5C-D**), indicating that maintaining free a fraction of free CP and RP may be important for proteostasis in stressed cells^8,25^.

### Assembly of TAADs is dependent on the cytoskeleton

While proteasome foci emerged almost instantaneously after hypertonic stress, the formation of TAADs spanned over several hours after adding aS aggregates to the media. To accelerate the internalization of aS aggregates by cells and validate the direct relationship between proteotoxicity and TAAD formation, we used a nanopipette for injection in PSMD14-eGFP cells. This is a bespoke approach enabling delivery of soluble aggregates at precise positions in the cytosol of a live cell^26^. Aggregates released into the cytosol induced TAAD formation within minutes (**Fig.2A**), while no response was observed when a buffer control or aS monomers at the same equivalent concentration were injected (**Fig.2B**). We therefore attribute the formation of TAADs to aggregate induction, rather than the presence of their constituent aS proteins.

**Figure 2.**
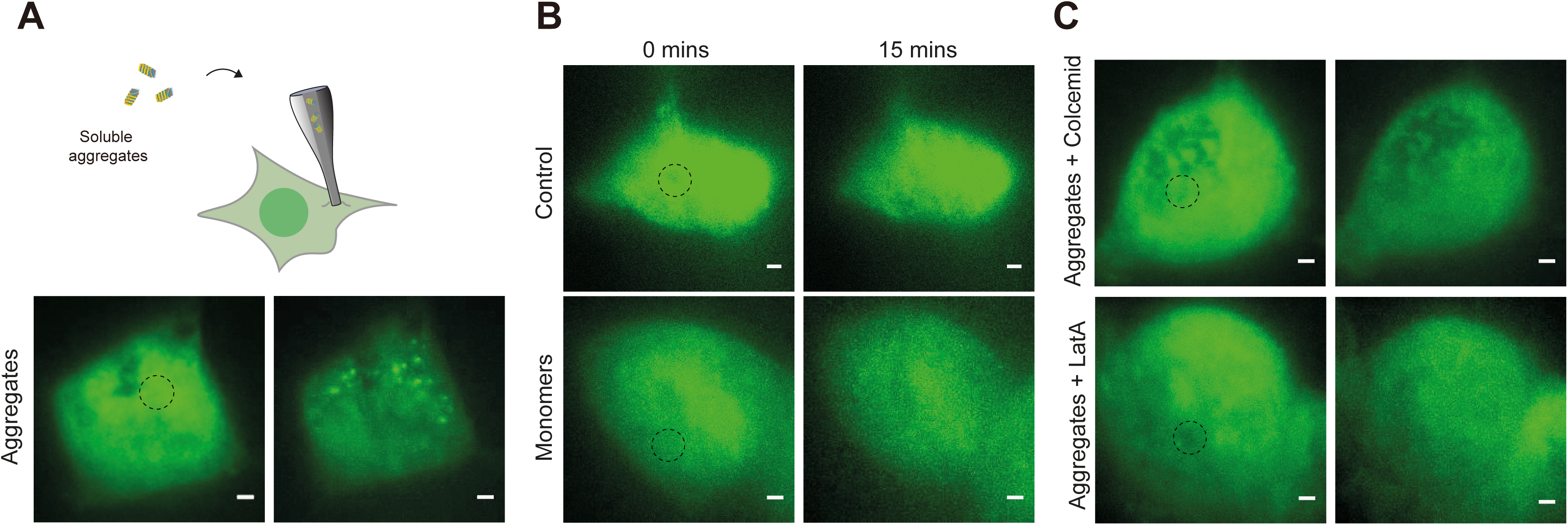
Proteasome response to aggregates in the cell is cytoskeleton-dependent (**A**) Schematic representation of aS aggregate injection into HEK293A cells expressing PSMD14-eGFP using a nanopipette, see **Materials and Methods**. (**B**) HILO images showing proteasomes forming foci in response to aS aggregates (see also **SVideo 1**) and compared to cells injected with aS monomers (**SVideo 2**). (**C**) No foci were observed in cells treated with 5 μM Colcemid or Latrunculin A (LatA) before injecting aggregates (see also **SVideo 3-4**), which disrupt the microtubule and actin networks, respectively. The injection positions are marked by circles with dotted lines.

We subsequently tested if TAAD formation was driven by active transport by through the cytoskeleton or facilitated through diffusion. Aggregate injection was repeated in PSMD14-eGFP cells pre-treated for 10 min with Colcemid or Latrunculin A (LatA), which depolymerizes microtubules or actin filaments, respectively (**Fig.2C**). No TAAD formation was observed under either condition, suggesting that interactions with the cytoskeleton is required to concentrate proteasomes into TAADs. Since hypertonic-induced foci were primarily localized in the nucleus, we thought their formation may be independent of the cytoskeleton. To test this, the cytoskeleton in PSMD14-eGFP and PSMB2-eGFP cells were depolymerized with Colcemid or LatA, respectively, prior to hypertonic stress. Hypertonic-induced foci appeared at similar levels to those observed in cells with intact cytoskeletal integrity, and remained present in the nucleus after 10 min (**Fig.S4C-D**), indicating that hypertonic stress-induced proteasome response is independent of the integrity of the cytoskeleton. This supports the notion that hypertonic-induced foci and TAADs differ in their mechanisms of formation.

### Cytoskeleton-dependent proteasomal translocation is linked to membrane potential

Invading aS aggregates are thought to interact and penetrate the cell membrane, which causes cell depolarization^27^, a phenomenon which has in turn been linked to diverting proteasome movements^28^. We therefore tested whether membrane potential (V_m_) could orchestrate proteasome transport inside PSMD14-eGFP cells, which would explain our observed TAAD formation. Performing single-cell patch clamp under TIRF microscopy, we found that hyperpolarization induced increased fluorescence intensity within the evanescent field of TIRF illumination closest to the coverslip, indicating proteasome accumulation close to the cell periphery (**Fig.3A-C** and **SFig.6A-D**). Expectedly, depolarization had the opposite effect and decreased the detected fluorescence intensity as proteasomes translocated into the cell interior. Oscillations in fluorescence intensity with voltage can be detected across the clamped cell, while the unclamped neighboring cells were unaffected (C1 and C2 cells, **Fig.3A** and **Fig.3C**). We did not notice morphological changes of the cell during patch clamp under our experimental conditions and a control protein of the ubiquitin system, eGFP-USP21^29^, did not respond to changes in V_m_ under the same experimental conditions (**Fig.3D** and **SFig.6E**). This suggests that the observed proteasome translocation is specifically related to membrane potential, toggling proteasomes towards cell interior or the plasma membrane through hyper- or depolarization, respectively.

**Figure 3.**
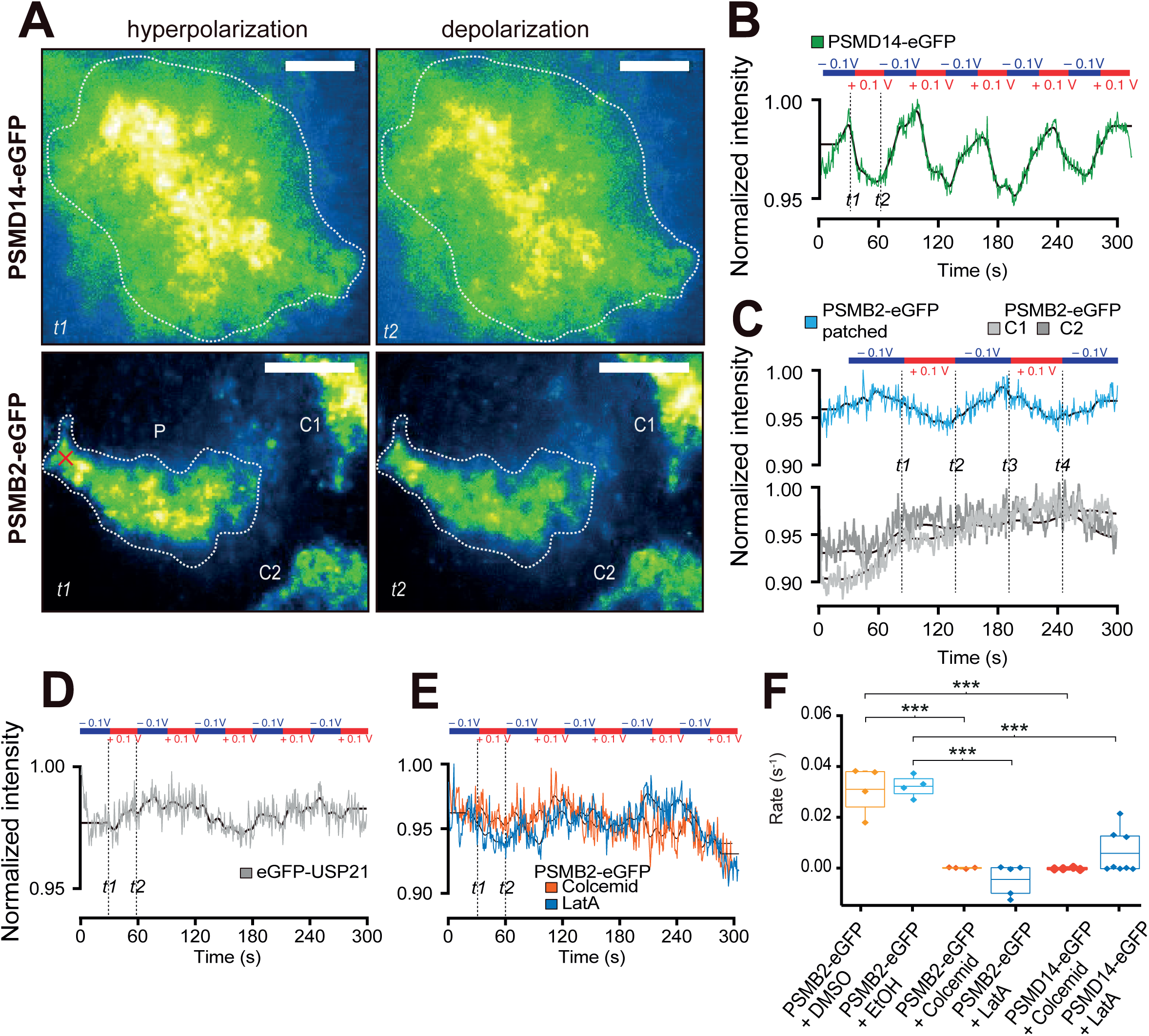
Membrane potential modulates proteasome localization. (**A**) Unpatched and patched PSMD14-eGFP (top) and PSMB2-eGFP (bottom) cells are imaged in TIRF mode with snapshots shown from various time points (four panels of *t1-t2* as in **B** and **C**), where the normalized fluorescence intensity is colored (see **Materials and Methods** for image processing and bleach correction). Three PSMB2-eGFP cells are visible in the bottom field of view, with only the cell on the left under patch clamp. The patch clamp position is marked with an “×” (red) on the cell (*P*) in panel *t1.* Regions of the patched cells used for intensity plots are marked in panel *t1* as dotted lines*. C1* and *C2* are used as control cells and their changes in intensity are shown in **C**. See also **SVideo 5**. (**B**) Changes in normalized fluorescence signal over time for the PSMD14-eGFP cell in **A**. Solid black lines represent signals after applying a filter in the frequency dimension to remove modulations smaller than the oscillation in potential. The clamped potentials applied are indicated on the top and oscillated between +100 mV (red) and −100 mV (blue) every 30 s. See also **SVideo 6**. (**C**) Changes in normalized fluorescence signal over time in PSMB2-eGFP cells for the patched cell and two control cells shown in **A**. (**D**) Cells expressing eGFP-USP21 did not show any change in fluorescence following changes in membrane potential induced by a whole cell patch-clamp (see also **SVideo 7**). (**E**) Depolymerization of microtubules and actin filaments affects potential-induced proteasome relocalization. PSMB2-eGFP cells were treated with 5 μM Colcemid or Latrunculin A (LatA) for 20 min at 37 °C and washed with cell media before mounted for TIRF imaging. PSMB2-eGFP cells treated with DMSO or ethanol (EtOH) vehicle controls showed similar results (see **SVideo 8**). Fluorescence intensity traces of the cells are plotted over time and the applied potential at each time indicated. (**F**) Repeated measurements of patch-clamp applied on PSMB2-eGFP cells treated with (*from left*) DMSO (mean rate with SD = 0.031 ± 0.009 nFI/s, n = 4), EtOH (0.032 ± 0.004 nFI/s, n = 4), Colcemid (0.0001 ± 0.0003 nFI/s, n = 4), LatA (−0.004 ± 0.006 nFI/s, n = 5). PSMD14-eGFP cells were also treated with Colcemid (−0.0002 ± 0.0002 nFI/s, n = 4) and LatA (0.006 ± 0.009 nFI/s, n = 8). All images are contrast-adjusted for optimal representation of the difference in intensity between hyper- and depolarized states.

To test whether such voltage-induced proteasome translocation is also cytoskeleton-dependent, we performed patch clamp on cells with compromised cytoskeleton. PSMB2-eGFP cells treated with Colcemid or LatA showed no proteasome translocation in response to hyper- or depolarization, while treatment with vehicle control (DMSO or EtOH) alone did not abolish potential-induced proteasome translocations (**Fig.3E-F**). Similar results were seen in PSMD14-eGFP cells treated with Colcemid or LatA (**Fig.3E-F**), confirming the dependence cytoskeleton in proteasome relocalization.

### Two distinct states of proteasomal movements

We subsequently examined the dynamics of proteasome motions at high time resolution by quantifying the movement of the individual particles in live cells using SPT. Tracking was achieved under TIRF mode and a statistical analysis of *Jump Distances* (JD) from individual tracks uncovered two diffusion coefficients (*D_fast_ and D_slow_*) (see **Materials and Methods**). The fast- and the slow-diffusing proteasome populations detected in PSMD14-eGFP cells were also observed in PSMB2-eGFP cells (**Fig.4A-B**). Since the ratio (1:1) and the diffusion coefficients (*D_fast_*at 3-4 µm/s^2^; *D_slow_* ∼0.5 µm/s^2^) of both populations were similar in PSMD14-eGFP and PSMB2-eGFP cells, we attribute this as evidence that common transport mechanisms are acting upon holoenzymes and individual RPs and CPs. Our findings are supported by a previous study in yeast proteasomes^30^, which assigned the fast population to freely diffusing particles and the slow population to interactions with macromolecular structures. Considering that TIRF imaging primarily detects proteasomes at the cell periphery, we validated diffusion coefficients and population ratios by repeating SPT in PSMB2-mEos or PSMD14-mEos cells and in HILO mode to image the cell interior (**SFig.8**), achieving similar results as in **Fig.4B**. We also performed SPT on eGFP-USP21, a component of the ubiquitin system similar in size to the free PSMD14 or PSMB2 but ∼40-fold smaller than the proteasome holoenzyme, and ∼16- and 12-fold smaller than the RP and CP, respectively. As the diffusion coefficients of eGFP-USP21 were significantly higher (**SFig.7**), we conclude that the eGFP and mEos-tagged PSMB2 and PSMD14 have been incorporated into proteasome particles, in agreement with **SFig.1B**.

**Figure 4.**
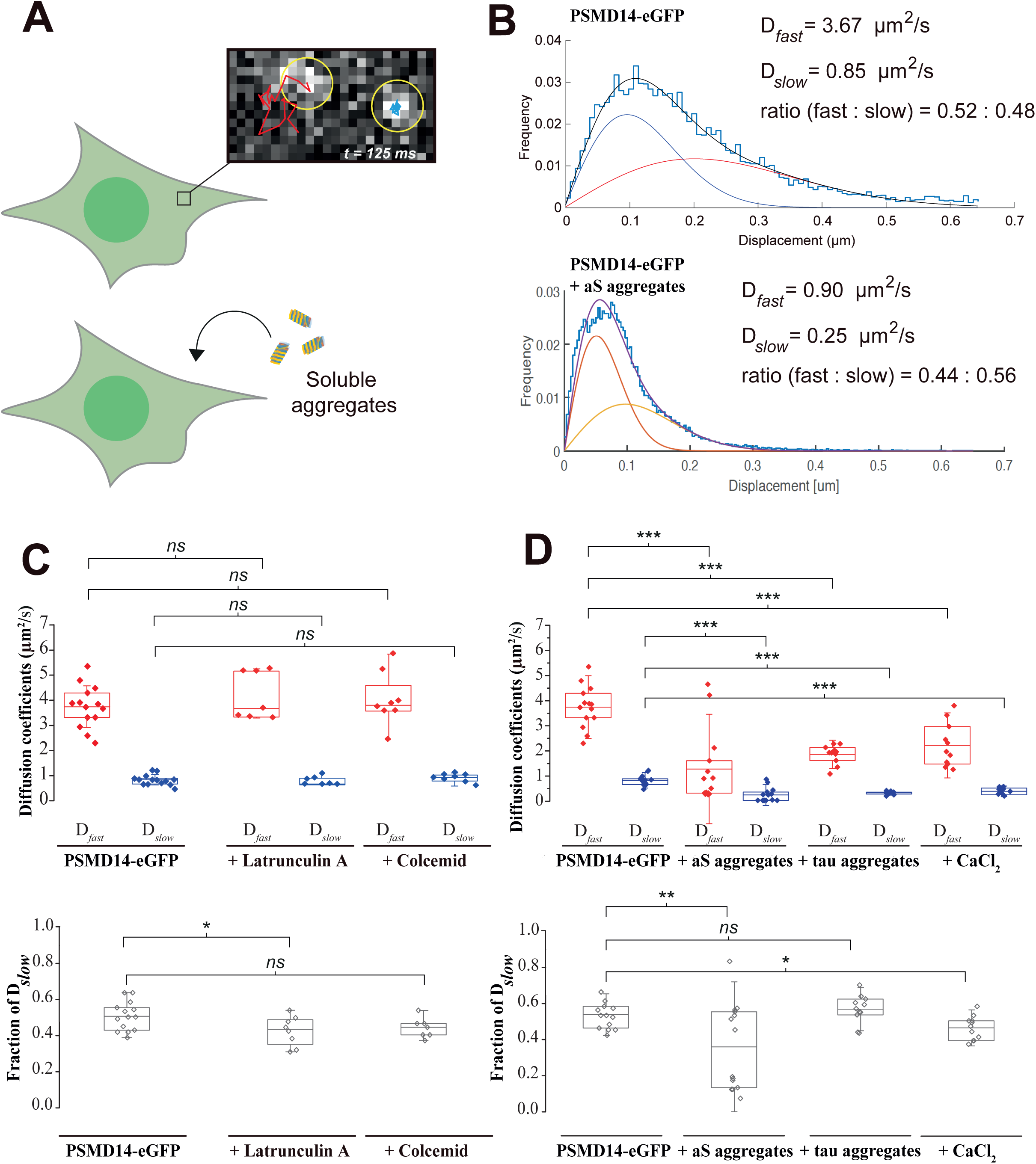
Single-proteasome tracking (SPT) by TIRF imaging in live cells shows two distinct proteasome behaviors. (**A**) A schematic of a proteasome moving fast (red tracks) inside a resting cell (top) expressing PSMD14-eGFP (pixel size = 107 nm, see also **SVideo 9**) and inside a cell incubated with aggregates (bottom). (**B**) A representative jump distance (JD) analysis (frequency of displacement distances in cyan bars) of all PSMD14-eGFP diffusion tracks detected within a cell (see also **SVideo 10**). Diffusion coefficients for the fast-(*D_fast_*, red line), the slow population (*D_slow_*, blue line) and overall fitting (black line) are shown with the ratio of *D_fast_* to *D_slow_* for this particular cell. Bottom JD frequency distribution shows a shift in the population distribution of the *D_fast_* and *D_slow_* proteasomes in response to aS aggregates. (**C**) Box plots quantifying the *D_fast_* and *D_slow_* (top) and the fraction of proteasomes in the fast population (bottom) from individual cells treated with Colcemid (n = 8) or LatA (n = 7) compared with control cells treated with DMSO (n = 14). (**D**) Similar box-plots describing cells at resting cells (n = 12) and cells treated with aS aggregates (n = 14), tau aggregates (n = 11), and CaCl_2_ (n = 11).

Since TAADs form in a cytoskeleton-dependent manner during proteotoxic stress (see **Fig.2**), we examined if the slow-diffusing population may be due to interactions with the cytoskeleton during resting state. Surprisingly, treating PSMD14-eGFP cells with Colcemid or LatA for 10 min prior to SPT did not alter *D_slow_* or *D_fast_*, nor did their population ratio change (**Fig.4C**). We also tested if the population of fast-moving proteasomes would reduce alongside TAAD formation. To examine this, we again incubated PSMD14-eGFP cells with aS aggregates and repeated the SPT approach as above. JD analysis still identified two diffusion populations following overnight aggregate incubation and with significantly decreased *D_fast_* and slightly increased ratio of the slow-diffusing population (**Fig.4D**). Together, this indicates a significant reduction in *D_fast_*, which reflects that proteasome diffusion is limited during TAAD but not hypertonic-induced foci formation. In support of stress-induced decrease in diffusion, proteasomes in cells under hypertonic stress incubated with CaCl_2_ also produced a similar change in their behavior, albeit with a less drastic reduction in the magnitude of *D_fast_* (**Fig.4D**).

### Proteasomes are globally confined as TAADs are formed

As aggregate clearance is a slow process^1^, we reasoned that proteasomes would be retained within the local vicinity in the form of TAADs to enable continuous disaggregation and degradation^31,32^. To demonstrate this, we computed the mean square displacement (*MSD*) from SPT of individual proteasomes and fitted them to three models, representing either freely diffusing particles, diffusive particles confined in a cage, or particles undergoing directive motion. Our data showed that the fractions of proteasomes that matched those models were in 3:4:3 ratio, respectively (**Fig.5A-C**). Comparing between resting and proteotoxicity-stressed conditions, we observed that cells expressing PSMD14-eGFP responded to aggregate incubation by increasing the fraction of RPs confined within molecular cages (**Fig.5D**). A smaller but significant increase in RPs traveling by active transport was also observed, while the free diffusing fraction reduced (**Fig.5D**). Similarly, PSMB2-eGFP cells also showed a significant increase in CPs traveling inside a confined volume due to proteotoxic stress, with a concomitant decrease in both the freely diffusive and active transported fractions (**Fig.5E**). Fitting these trajectories to model the radius of confinement suggests proteasomes are caged within a volume with an approximate radius of 0.39±0.04 μm (see **Materials and Methods**). Interestingly, αS aggregates decrease the size of confinement in stressed cells when compared to confined proteasomes in resting cells, but not tau (**SFig.9**). This difference in radius may be a consequence of the distinct aggregate size and morphology associated with each type of protein aggregate, or that different cellular enzymes are recruited to target aS and tau aggregates. Taken together, our data suggest a model where the cellular response to aggregate invasion involves a tightly organized redistribution and trapping of proteasomes to address the challenges of proteotoxic stress (**Fig.5F-G**).

**Figure 5.**
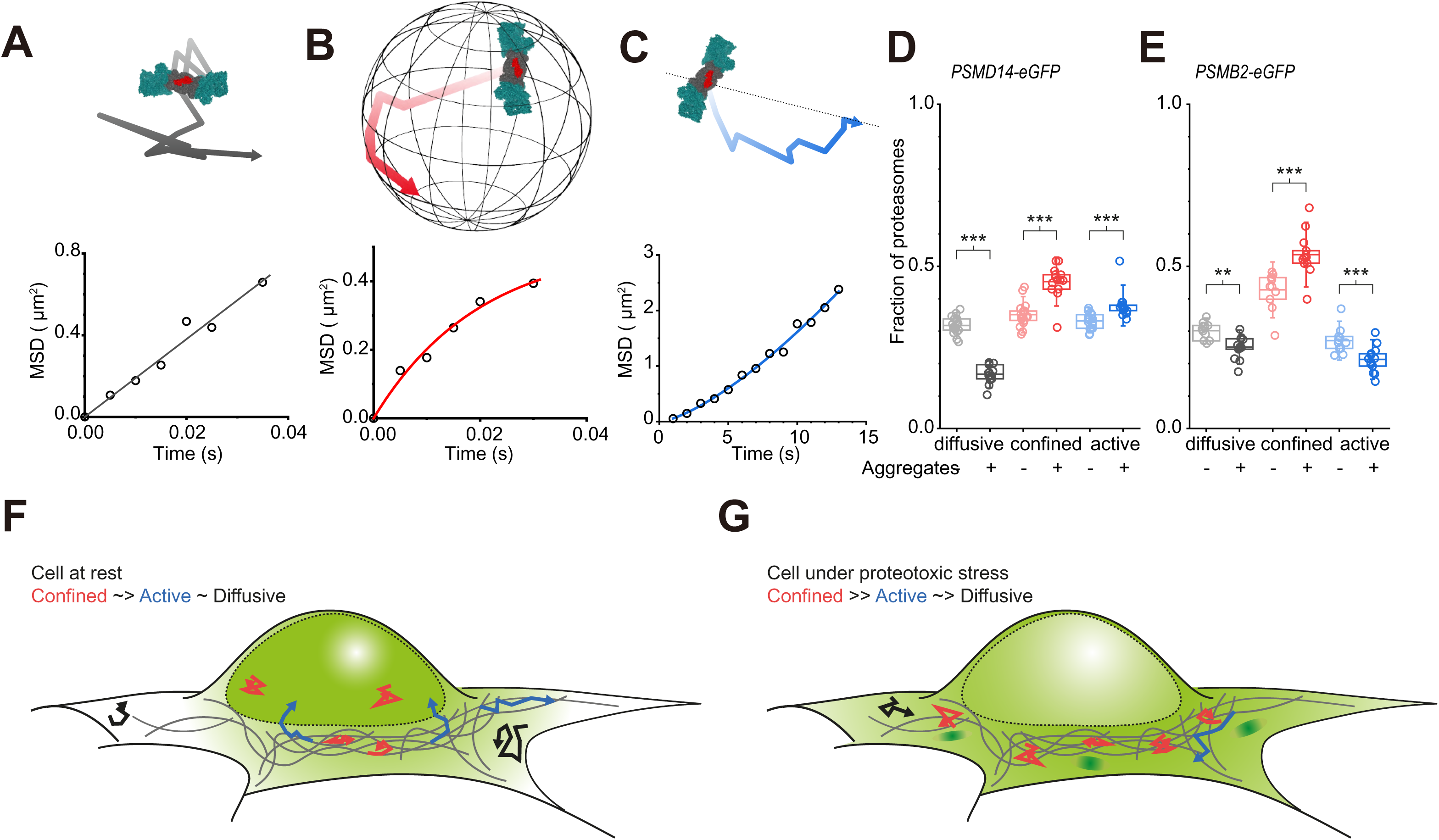
Mean squared displacement (MSD) analysis of SPT data. Individual tracks recorded from PSMB2-eGFP or PSMD14-eGFP molecules were classified as (**A**) stochastic diffusion, (**B**) confined diffusion, and (**C**) active transport. (**D**) The fraction of molecules in each cell that was best described as diffusion, confined diffusion, or active transport, are plotted from resting cells expressing PSMD14-eGFP (n = 18 cells), and cells incubated with aS aggregates (n = 14). (**E**) Similarly, resting cells expressing PSMB2-eGFP (n = 11) and after incubation with aS aggregates (n = 10) were also characterized by MSD analysis. (**F**) Schematic comparing the global proteasome redistribution of cells at rest, and (**G**) under proteotoxic stress. Proteasomes in resting cells are predominantly localized to the nucleus (green shading). Cells under proteotoxic stress rebalance their proteasome localization, and can form foci at gradation centers (dark green shading). Individual proteasomes are increasingly confined (red tracks) in cells under proteotoxic stress compared to resting cells, while the fraction of proteasomes which can freely diffuse (black tracks) are significantly reduced. Active transport (blue tracks) of RP and CP show slight increases and decreases respectively induced by proteotoxic stress, potentially indicative of the energy costs involved in redistributing the respective free subunits against concentration gradients.

## Discussion

While the diffusional and active transport elements of proteasomes have been reported previously, we demonstrated in this study how proteasomes respond to stress by altering their mode of motions to effectively increase proteasome concentration at sites of degradation. This would enable increased degradation activity, e.g. against protein aggregates. Our findings suggest that the distribution and motion of CPs and RPs were close to identical and could not be quantitatively distinguished by localization or re-organization induced by stress. It is therefore very likely that proteasome holoenzymes remain assembled at a stable level in equilibrium with free CP and RPs.

Slow-diffusing proteasomes were previously attributed to possible interactions with other large macromolecules^30^. We further expand this view and suggest that under changing cellular conditions, proteasomes may also associate reversibly with cellular organelles^11,33,34^ or accumulate together to form cellular bodies, which could account for the slow-diffusing population. Our results further show that the balance between the three modes of proteasome motion: confined- and free diffusion and active transport is carefully modulated and tuned to specific cellular requirements. This enables proteasomes to reorganize upon proteotoxic stress, increasing the proportion of proteasomes confined in TAADs and at other centers of degradation^19,20,35,36^. By allowing reorganization of proteasomes, it follows that the availability of degradation activity is also redistributed during aggregate clearance, thus likely deprioritizing normal cellular proteostasis activities. Possibly, delayed return of confined proteasomes to other subcellular regions may contribute to the decline in proteostasis. This may become further aggravated should aggregates become degradation-resistant and prolong proteasome confinement, thus impeding proteostasis events.

Similarly, the observed fraction of freely diffusing proteasomes under resting state may also be by design to minimize time and energy redistributing proteasomes to nascent degradation centers, such as TAADs. For instance, stalling proteasomes from a reservoir of freely diffusing proteins may be more efficient than actively transporting proteasomes from distal sites. The observed changes in the proportion of proteasomes under active transport are relatively small compared to the changes in the fractions of freely diffusing and confined proteasomes. This may reflect how the cell only consumes energy moving proteins when it must direct them against concentration gradients to fulfil critical proteostasis needs at specific subcellular locations requiring greater concentration of proteasomes. Since the turnover of many proteins are controlled by degradation, small changes in local proteasome levels may be sufficient to activate physiological processes sensitive to degradation. It remains to be discovered in future studies how distinct processes associated with or regulated by proteasomal degradation may be affected by altering their distribution.

## Acknowledgement

MJM, YZ, BL, LS, JEL, JXM, JK, KDPP and YY performed experiments and analyzed the data. AHL, MB, AP, ARC provided reagents, experimental setup and support. YY, DK, DF, SFL guided experimental design. YY conceptualized the research and prepared the manuscript with MJM. This work was funded by a Sir Henry Wellcome Research Fellowship [101585/Z/13/Z] awarded to YY. Work in YY lab is funded through an ARUK Major Projects Grant [ARUK-PG2023A-025].

## Supplementary Figures

**SFigure 1.** Genomic expression of CRISPR-edited and FP-tagged PSMB2 or PSMD14 subunits are incorporated into proteasome particles. (**A**) Molecular structure of the mammalian 26S proteasome holoenzyme (pdb-id: 5GJQ) highlighting the protease subunits responsible for peptidase activities of the 20S core particle (CP, in shades of gray) and the ATPases providing substrate unfolding and translocation activities of the 19S regulatory particles (RPs, in shades of cyan). (**B**) PSMB2 (red) and PSMD14 subunits are highlighted within the CP and RP, respectively, of the proteasome. Lysates from CRISPR-edited cell lines expressing (**C**) PSMB2-eGFP or PSMD14-eGFP, and (**D**) PSMB2-mEos or PSMD14-mEos are separated by native gradient-gel (4-16% acrylamide) electrophoresis and visualized by fluorescence excitation at 488 nm (*left*) for eGFP or mEos detection. Gels were incubated with 100 µM of LLVY-AMC, a fluorogenic proteasome substrate, and visualized by excitation at 350 nm (*right*). Purified 26S proteasome holoenzyme was loaded as a control and the position of free CP and RP and singly- and doubly-capped holoenzymes are indicated. Molecular weights markers are shown to demonstrate the absence of any free GFP-tagged subunits not incorporated into the proteasome.

**SFigure 2.** Determination of the percentage of endogenous proteasome subunits labeled by FP. Western blots against (**A**) PSMB2 and (**B**) PSMD14 in three biological repeats each of cell lines expressing PSMB2-eGFP, PSMB2-mEos, PSMD14-eGFP or PSMD14-mEos. (**C**) Percentage of labeled proteasome subunits was then determined from densitometry analysis of the FP-modified band as a percentage of total (FP-modified+unmodified), averaging from three repeats rounded to the nearest integer.

**SFigure 3.** Light-sheet imaging validates that proteasome distribution changes upon aS aggregate invasion. (**A**) Light-sheet imaging of PSMD14-eGFP cells at rest (left) or treated with recombinant aS aggregates, performed using a bespoke setup previously described^37^. The change in distribution and formation of proteasome foci are evident from the 3D image, shown as orthogonal views. (**B**) Nanopipette injection of Alexa647-labeled aS aggregates demonstrates colocalization of proteasome foci (green) and aggregates (red) inside cells. See **Materials and Methods** for details.

**SFigure 4.** The formation of proteasome foci induced by hyperosmotic stress is not cytoskeleton-dependent. (**A**) Untreated PSMB2-eGFP or PSMD14-eGFP cells were compared to (**B**) cells incubated with 150 mM CaCl_2_ for 10 minutes. (**C-D**) The formation of the proteasome foci were not prevented by pre-incubation of cells with 5 μM Colcemid or LatA prior to the addition of CaCl_2_.

**SFigure 5.** Immunofluorescence staining of differentiated SH-SY5Y cells shows free CP and RP in the cell and in complex. (**A)** Untreated SH-SY5Y cells were fixed and stained with antibodies to label CP (PSMA2, red) and RP (PSMD14, green). Insets show foci depicting singly- and doubly-capped proteasomes. Scale bars: 5 μm, and insets 50 nm. (**B**) Colocalization of CP (left) and RP (right) enabled calculation of uncapped versus singly- and doubly-capped proteasome ratios in resting cells. (**C**) CP and (**D**) RP box plots show that incubation of 1 μM aS aggregates do not induce any significance changes between the relative ratios of free particles or holoenzymes in the cytoplasm. (n = 20 resting cells and n = 21 cells incubated with aS aggregates).

**SFigure 6.** Current(I)-potential(V) relationship from patch-clamped HEK cells imaged by TIRF microscopy. (**A**) A typical set of voltage responses to current injections in a PSMB2-eGFP cell and (**B**) corresponding IV curve of a recorded cell. (**C**) Voltage control (+100 mV or -100 mV) of recorded PSMB2-eGFP cells with 30 s step plotted here against the current response to the voltage control. (**D**) Averaged current response range of HEK cells at holding potential +100mV or -100 mV (n = 14 cells). (**E**) Rate of change of normalized fluorescence intensity of cells expressing eGFP-USP21 with respect to cycling between depolarizing and hyperpolarizing membrane potentials, compared to PSMB2-eGFP and PSMD14-eGFP cells.

**SFigure 7.** Single-particle tracking of eGFP-USP21 expressed in HEK cells (n = 8 cells) and used as a control protein of similar molecular weight as PSMD14 but does not form part of the proteasome, presented as in **Fig.4**.

**SFigure 8.** Single-particle tracking of PSMD14-mEos and PSMB2-mEos cells. (**A**) TIRF imaging of (*left*) PSMD14-mEos cells (mean ± SD for *D_fast_* = 4.1 ± 1.1 and *D_slow_* = 0.9 ± 0.3 µm^2^/s, relative abundance of *D_fast_* 0.52 ± 0.1, n = 10 cells) and (*right*) Pre1-mEos cells (*D_fast_* = 3.05 ± 0.8 and *D_slow_* = 0.7 ± 0.2 µm^2^/s, where *D_fast_* represents 0.51 ± 0.1, n = 8 cells). (**B**) Tracking in HILO mode (n = 13 and n = 16 cells, respectively, of PSMD14-mEos and Pre1-mEos cells) gives similar diffusion coefficients as in **Fig.4**, suggesting that proteasomal particles diffuse with similar behavior across the whole cell. Due to increased photo-bleaching in HILO mode, tracks from multiple cells were combined for jump distance analysis.

**SFigure 9.** Proteasomes undergoing confined diffusion were identified from cells expressing PSMB2-eGFP during single-particle tracking experiments. MSD curves calculated from the single-particle tracks of these proteasomes were fitted to quantify the radius of confinement of cells treated with tau (n = 14) and aS (n = 10) aggregates for 12 hrs compared with resting cells (n =11).

## Supplementary Videos

Supplementary Videos can be found on the online version of this manuscript and on youtube.com: (https://www.youtube.com/playlist?list=PLgc4yUNxF7D21SY7J0HPBDZCoGrzsq0RR)

**SVideo 1.** Recombinant aS aggregates were injected into the cytosol of a PSMD14-eGFP cell and recorded at 33 ms frame rate in HILO mode.

**SVideo 2.** Recombinant aS monomers were injected into the cytosol of a PSMD14-eGFP cell and recorded at 33 ms frame rate in HILO mode.

**SVideo 3.** Recombinant aS aggregates were injected into the cytosol of a PSMD14-eGFP cell pre-treated with colcemid for 10 min and recorded at 33 ms frame rate in HILO mode.

**SVideo 4.** Recombinant aS aggregates were injected into the cytosol of a PSMD14-eGFP cell pre-treated with cytochalasin D for 10 min and recorded at 33 ms frame rate in HILO mode.

**SVideo 5.** Patched PSMB2-eGFP cell (*left*) and control cells (*right*, top and bottom) imaged in TIRF mode undergoing oscillation between hyperpolarization (-100 mV, blue) and depolarization (+100 mV, red) recorded at 60 ms per frame. Scale bar indicates 5 µm. The video was contrast-adjusted for optimal representation of the difference in intensity between hyper- and depolarized states.

**SVideo 6.** Patched PSMD14-eGFP cell imaged in TIRF mode undergoing oscillation between hyperpolarization (-100 mV, blue) and depolarization (+100 mV, red). Scale bar indicates 5 µm. The video was recorded at 60 ms per frame and contrast-adjusted for optimal representation of the difference in intensity between hyper- and depolarized states.

**SVideo 7.** Patched cell expressing eGFP-USP21 from a plasmid vector and imaged in TIRF mode undergoing oscillation between hyperpolarization (-100 mV, blue) and depolarization (+100 mV, red). Scale bar indicates 5 µm. The video was recorded at 60 ms per frame and contrast-adjusted for optimal representation of the difference in intensity between hyper- and depolarized states.

**SVideo 8.** Patched PSMB2-eGFP cells treated with Colcemid (*top left*), LatA (*top right*), DMSO (*bottom left,* reduced to 62.5% of original size) or ethanol (*bottom right*) imaged in TIRF mode undergoing oscillation between hyperpolarization (-100 mV, blue) and depolarization (+100 mV, red). Scale bars indicate 5 µm. All videos were recorded at 60 ms per frame and contrast-adjusted for optimal representation of the difference in intensity between hyper- and depolarized states.

**SVideo 9.** TIRF imaging of two PSMD14-eGFP molecules with distinct movement pattern, as shown in **Figure 4A**. Scale bar indicates 0.5 μm.

**SVideo 10.** TIRF imaging of PSMD14-eGFP (***A***) and PSMB2-eGFP (***B***) diffusion in live cells recorded at 5 ms per frame. Scale bars indicate 2 µm.

## Materials and Methods

### Transgenic cell lines

Genomic sequences of mammalian *PSMD14* and *PSMB2* were modified at 3’-end of the last exon with CRISPR-cas9 genetic engineering system to replace the stop codon with DNA sequences coding for eGFP or mEos3.2. HEK293A (HEK) cells were co-transfected with a donor plasmid (encoding the FP sequence flanked by the homology arms of *PSMD14* or *PSMB2*), gRNA plasmid (targeting the 3’-end of either *PSMD14* or *PSMB2* DNA sequence), and an SpCas9 expression plasmid (Addgene #62988). Clones expressing eGFP or mEos were individually sorted by fluorescence intensity and grown to confluency. Correct insertions of FP sequence into the genome were identified by PCR using primers based on the flanking sequences and confirmed by sequencing. Cell lines deemed suitable for TIRF imaging were selected for subsequent experiments. All cell lines used in this study are clonal and heterozygous at the modified locus, expressing both unmodified and FP-modified proteasomal subunits. The FP was detected at a suitable level for single-molecule imaging in cells.

### Western blot analysis of cell lysates

PSMD14-eGFP, PSMD14-mEos, PSMB2-eGFP, and PSMB2-mEos knock-in cells were washed thrice in PBS, and collected by scraping in lysis buffer (50 mM Tris pH 7.4 at 4°C, 5 mM MgCl_2_, 2 mM ATP). Cells were lysed by homogenization using a motorized pestle mixer (431-0095, VWR) for 30 s and centrifuged on a benchtop centrifuge at 21,000 × g for 20 min at 4°C. The supernatant was transferred to new tubes, and the concentration measured by Bradford assay. Cell lysates (20 µg) were resolved on gels and transferred to 0.45 µm PVDF membranes (Bio-Rad) using Trans-Blot semi-dry Transfer System at 25 V, 2.5 A for 15 min. Membranes were then blocked in 5% BSA in Tris-buffered Saline with 0.05% Tween 20 (TBS-T) for 1 hr at 4°C, and incubated with primary antibody for PSMD14 (MA5-35818, Thermo Fisher) or PSMB2 (sc-58410, Santa Cruz) overnight at 1:1000 dilution. Following three washes for 5 min in TBS-T (20 mM Tris pH 7.6 at room temperature (RT, 22°C), 150 mM NaCl, 0.1% Tween 20), membranes were then incubated with secondary antibodies Goat anti-rabbit Alexa Fluor 488 (A-11008, Thermo Fisher) or Goat anti-mouse Alexa Fluor 647 (A-21235, Thermo Fisher) at 1:10,000 dilution for 1 hr at RT. Finally, membranes were washed thrice for 10 min in TBS-T before fluorescence of secondary antibodies was detected on an iBright imaging system (Invitrogen).

### SH-SY5Y maintenance and differentiation

SH-SY5Y human neuroblastoma cells were maintained in Dulbecco’s modified Eagle’s medium (DMEM), supplemented with 10% (v/v) fetal bovine serum (FBS), at 37°C in 5% CO_2_, and passaged using 0.05% Trypsin-EDTA. To differentiate SH-SY5Y cells, an established protocol was used^38^. Briefly, cells were plated at a density of 10 k/cm^2^ on coverslips. The following day media was changed to DM1 (DMEM (no sodium pyruvate) supplemented with 5% FBS and retinoic acid (10 μM)) every day for three days. Cell media was changed to DM2 (Neurobasal A, supplemented with N2 (1X), L-glutamine (1 mM), and BDNF (50 13 ng/μl)) every day for a further three days.

### Alpha-synuclein and tau aggregation

Recombinant aS and full-length tau P301S mutant (isoform 0N4R) were purified following protocols as previously described before^31,32,39^. Aggregation of αS was completed at 70 µM in reaction buffer containing 50 mM Tris, 150 mM NaCl, pH 7.4, 0.1% NaN_3_ incubated at 37□°C, shaking at 200 rpm for 72 hrs. Tau aggregates were formed using 2 µM tau monomers in PBS incubated with an equimolar final concentration of low-molecular-weight heparin (average 5 kDa; Thermo Fisher Scientific) at 37°C for 24 hrs without shaking.

### Cell imaging preparations

Cells were plated on glass coverslips (0.17 mm, Thorlabs) and grown at 37°C in DMEM (Sigma) containing 10% FBS and 1% penicillin-streptomycin one day before imaging. All glass coverslips were cleaned with Argon plasma for 1 hr before incubation with cells. For SMLM imaging of mEos localizations, fixed cells were incubated with PBS buffer containing 4% paraformaldehyde (PFA) and 0.2% glutaraldehyde for 10 min on the bench. Prior to imaging, cells were washed thrice and imaged in warm OptiMEM (Sigma) pre-filtered through 0.02 µm pores. A metal chamber was used to secure the coverslip above the objective. All imaging was performed in temperature-controlled chambers. For microtubule or actin depolymerization assays, cells were incubated with 5 µM Colcemid (Sigma) or Latrunculin A (Anachem), prepared in DMSO or ethanol, respectively. After 10 min at 37°C, cells were taken out from the incubator and washed three times using OptiMEM before imaging as described above.

### Light-sheet imaging

Cells were imaged using single objective cantilever SPIM (socSPIM), as detailed previously^40^. Briefly, a commercial AFM cantilever (ContAl-G, BudgetSensors) above the objective of an inverted microscope (Eclipse TiU, Nikon). The cantilever was coated with reflective aluminium, and was fitted onto a machined brass rod using cyanoacrylate adhesive. A cylindrical lens placed before the cantilever generated a light-sheet with an axial thickness of ∼1.9 μm FWHM. The sample stage was moved by a xyz piezo (P-611.3 Nanocube, Physik Instrumente) allowing the light-sheet to scan through the sample at 30 ms exposure time with scanning steps of 200 nm. Scanning data was assembled into hyperstacks and rendered using the 3D projection plugin and 3D viewer plugin in ImageJ/FIJI.

### Immunofluorescence labeling to assess relative fractions of singly- or doubly-capped proteasomes

Cells were fixed with 4% PFA in PBS (20 min at RT). Cells were washed thrice in PBS and permeabilization was performed with Triton (0.1%, in PBS, 15 min at RT). Blocking was performed for 1 hr at RT with 10% (v/v) goat serum (GS) in PBS. Primary antibody incubation was performed in 5% GS overnight at 4 °C. After three washes with PBS, secondary antibody incubation (5% GS in PBS) was performed for 1 h at RT, protected from light. Cells were washed three times with PBS prior to imaging. Primary antibodies against RP (rabbit anti-PSMD14, 1:400) and CP (mouse MCP21, 1:400) and secondary anti-rabbit-Alexa568 (1:1000) and anti-mouse-Alexa488 (1:5000) were used. Monomers of aS were also labeled with Alexa647, and were aggregated with unlabeled monomers at a 1:9 ratio labeled:unlabeled under shaking conditions and sonicated, as previously described, to produce labeled alpha-synuclein aggregates capable of passing through cell membranes^1^. Cells were incubated with 1 μM labeled aggregates for 24 hrs before cells were imaged.

### SMLM imaging of immunofluorescence labeled cells

Cells were imaged using an ECLIPSE Ti2-E inverted microscope (Nikon), coupled to a sCMOS camera (Prime95B, Photometrics) and C-FLEX laser combiner (HÜBNER Photonics), as previously described^1^. Briefly, imaging was completed with fixed cells incubated with imaging buffer (PBS with 1□mg/mL glucose oxidase, 0.02□mg/mL catalase, 10% (w/w) glucose, and 100□mM methylamine). Each field of view was recorded as a set of 2000 frame movies sequentially for each fluorophore. Multi-color images of each field of view were reconstructed in to a single frame using Matlab scripts^41^. Bursts of fluorescence from individual molecules were identified after applying a box filter and thresholds for each pixel, which were used based on that specific pixel’s variance when measuring background noise. Single-molecule localisations from each frame were calculated by fitting the point spread function to a 2D Gaussian profile. Chromatic aberrations were corrected using a second order polynomial function, as determined by imaging Tetraspeck beads (0.1 µm, fluorescent blue/green/orange/dark red; Life Technologies) across the emission wavelengths used in the experiment. Localizations from all frames and colors were aligned and plotted to produce images with resolutions of approximately 25 nm^1^.

### HILO 3D imaging of cells

Cells were imaged using the ECLIPSE Ti2-E inverted microscope (Nikon) described above. The imaging buffer for observing live cells was Fluorobrite supplemented with 10% FBS. The E-TIRF arm was used to position the laser beam to illuminate the sample at a angle suitable for HILO imaging. The objective was moved axially in steps of 100 nm to image the entire cell volume, and 3D representations of each cell were rendered using the 3D viewer Image/FIJI plugin.

### Image analysis to calculate fractions of singly- and doubly-capped proteasomes

Regions of interest were identified within multi-color reconstructed SMLM images of cells labeled with fluorescent antibodies against CPs and RPs, respectively. Each region of interest was analyzed using the ImageJ/FIJI plugin ComDet (v.0.5.5). Holoenzymes were identified as CP and RP clusters within 100 nm of each other. To determine the number of RPs in each holoenzyme, the RP fluorescence intensities for each cluster in that cell were used to calculate the average intensity of a single RP. The number of singly- and doubly-capped holoenzymes were calculated and are reported as fractions of the total number of CP or RP clusters, respectively.

### Nanopipette delivery and imaging of aS aggregates

Nanopipettes were fabricated from quartz capillaries to achieve ∼300-400 nm inner and ∼600-800 nm outer diameter with 40-80 MΩ electronic resistance, detailed elsewhere^26^. Briefly, the nanopipette was backfilled with 8 μl of 1.4 μM aS in PBS, with an Ag/AgCl electrode inserted inside. The pipette was immersed into a bath of PBS buffer, which contained cell samples, while another Ag/AgCl electrode served as the reference electrode was put into the buffer. A +100 mV voltage bias was applied between the electrodes inside the nanopipette and the bath, generating a +2 nA ionic current. Prior to injection, the nanopipette was first moved down to a position ∼15 μm above the cell surface using the white light microscope and the piezo moved the nanopipette down to the target area at a speed of 25 μm/s, while the ion current was recorded in field programmable gate array (FPGA) at the same time. Once the nanopipette had approached the cell surface, the piezo moved the pipette down by an extra 5 μm in one step to penetrate the cell membrane. Once the nanopipette had penetrated the cell membrane, a voltage pulse (-500 mV, 10 s) was applied to force the molecules out of the nanopipette and into the cytoplasm. After the injection process was finished, the nanopipette was retracted back by 5 μm in one step preventing the penetrated cell sticking on the nanopipette and moving up with it.

### Imaging in TIRF and HILO for single-particle tracking

Imaging of live cells was performed using a bespoke TIRF microscope. Diode lasers operating at 405 nm (Cobolt MLD), 488 nm (Toptica) or 561 nm (Cobolt Jive 500) were directed into an oil-immersion TIRF objective (60× Apo N TIRF, NA 1.49, Olympus) mounted on an Eclipse Ti2 microscope (Nikon Corporation). The change from HILO to TIRF mode was achieved by altering the beam offset perpendicular to the optical axis resulting in TIRF/HILO imaging. Emitted fluorescence was collected by the same objective and separated from the returning TIR beam by a dichroic filter (Di01-R405/488/561/635-25×36; Semrock). Emission filters were applied for eGFP (FF01-496/LP-25, FF01-520/44-25) or mEos (BLP02-561R-25, FF01-607/36-25) imaging. Images were recorded on an EMCCD camera (Evolve 512Delta, Photometrics), by alternating between laser excitation sources (561 nm laser at 0.8 kWcm^-2^, 488 nm at 1.5 kWcm^-2^ and 405 nm at 0.7 kWcm^-2^, determined by measuring the power exiting the objective and the footprint of the beam in the imaging plane) using mechanical shutters (Prior Scientific) and an exposure of 55 ms (localization). For single-molecule tracking, the measured exposure time was 5 ms. A short pulse of 405 nm was used to photoconvert mEos, which was subsequently excited using the 561 nm laser. The pixel size (107 nm) was measured prior to recordings.

### Single-molecule tracking analysis

Single-molecule images were analyzed using a custom Matlab script^42^. A bandpass filter (1 to 5 pixels) was first applied, followed by identification of local maxima in raw images. Maxima were identified based on a signal-to-noise threshold, which was set to 4, and spots with a radius less than 8 pixels were then linked together to form tracks^43^. The tracking algorithm allowed molecules to move a maximum distance of 6 pixels (8 pixels for eGFP-USP21) per frame (5 ms). Only tracks longer than 6 frames were used for analysis.

### Diffusion analysis by jump distance

Jump distance (*JD*) analysis has been described before^37,42^ and was applied to tracks detected from each imaged cell to investigate heterogeneities in the diffusion coefficient distribution.

Briefly, the probability distribution *P*(*r2*, Δ*t*) describing the probability of a particle travelling a distance between *r* and *dr*, t in one time step, *Δt*, was fitted using **Equation 1**.:

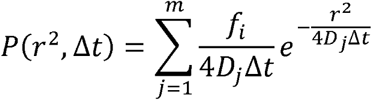

where *f* is the fraction corresponding to population *j*, and *m* is the number of populations. *JD* analysis was applied to tracks from each cell individually. *m* was selected by evaluating the coefficient of determination, *R^2^*, of the fit to a single population. *m=1 for R^2^>0.9* and *m=2 for R^2^<=0.9* (detailed in reference ^42^).

### Diffusion analysis by mean square displacement

The MSD curve for each single particle track was calculated using Kehl^44^, a Matlab function written to calculate MSD plots avoiding using loops. MSD curves were calculated from SPT tracks using the sub-pixel localisation of particles at time *t* and *t*+ *τ*. All displacements with the same value of *τ* are used to calculate a mean square displacement, and a MSD plot is recorded as a function of time intervals, *τ*, as described by **Equation 2**.:

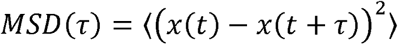

Each particle’s behaviour was classified as either freely diffusing, diffusing in a cage, or being actively transported according to methods based on Otero et al.^33^. Briefly, the MSD plot is fitted using **Equation 3**.:

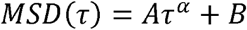

where *α* can vary between 0 and 2. Particles were described as freely diffusing if *α*< 0.5, confined in a cage if 0.9<*α* < 1.1, or being actively transported if *α* > 1.25 (particles were discarded if *α* lies outside these ranges). The radius of confinement was calculated by fitting each MSD curve classified as a confined particle to **Equation 4**.:

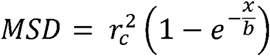

where 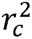 represents the plateau of the MSD curve, and the radius of confinement (*r_c_*) for each particle. The mean radius of confinement for all particles each cell is plotted in **Figure S9**.

### Patch clamp and imaging

Recording pipettes (4–6 MΩ) were pulled from borosilicate glass, and were filled with an internal solution (135 mM potassium gluconate, 10 mM HEPES, 10 mM sodium phosphocreatine, 4 mM KCl, 4 mM MgATP, 0.3 mM Na2GTP, pH 7.2 [adjusted with KOH]) and osmolarity set to 291 mOsmol l^−m^. Signals were acquired using an Axon Multiclamp 200B amplifier (Molecular Devices). Recordings were low-pass filtered at 2 kHz and acquired at 5 kHz using an NI X Series Multifunction DAQ (National Instrument) programmed by Axograph software. Membrane voltage was recorded in the current clamp mode. Series resistance was 10–25 Mso and experiments were terminated if this range was exceeded.

To investigate proteasomal responses to membrane potential, the membrane potential of cells was manipulated in voltage clamp mode with a 30 s step (unless otherwise specified) of -100 mV or +100 mV alternatively. Current traces were filtered at 2 kHz and digitally sampled at 20 kHz using AxoGraph software. The bath level of the recording chamber was kept to a minimum to reduce pipette capacitance. Leak current was subtracted off-line using a template constructed from the currents induced by a -10 mV hyperpolarizing step. The resting V_m_ was found to be around -20 mV, consistent with published values for HEK cells (e.g. [*(44, 45)*]). V_m_ was then clamped on cells expressing PSMB2-eGFP or PSMD14 and oscillated between depolarization (+100 mV) and hyperpolarization (-100 mV). A lower laser power was used for TIRF illumination to reduce the rate of eGFP photo-bleaching during patch clamp experiments.

White-light transmission and fluorescence images were acquired with a modified fluorescent microscope based on an inverted microscope, equipped with an infinity-corrected oil immersion objective (Olympus UPlanApo, 100×, NA 1.4), operating in TIRF imaging mode to reduce the excitation volume, and detected on a 512 × 512 pixel EMCCD (Evolve 512Delta, Photometrics) at a rate of 60 ms per frame for proteasome imaging. The filters used were a dichroic mirror (Di01-R405/488/561/635-25x36, Semrock) and a 525/50 nm band pass filter (FF03-525/50-25; Semrock). Laser excitation was provided by a 488 nm solid-state laser (Spectra-Physics) at a power density of ∼0.7 kWcm^-2^.

### Potential regulated imaging analysis

Image Analysis was done using Fiji and custom-written Matlab code. Image stacks were bleach-corrected using the in-built exponential *bleach correction* function of Fiji (Kota Miura et al. (2014). ImageJ Plugin CorrectBleach V2.0.2. Zenodo. 10.5281/zenodo.30769) applied to the cell of interest. Subsequently, the stacks were 10-fold binned in time to generate frame rates of 600 ms. For the image analysis time-traces from the cell of interest were low-pass filtered at a pass band frequency of 0.05 Hz, a steepness of 0.85 and stop band attenuation of 30 dB. The resulting traces were then normalized to their maximum intensities and analyzed using the Matlab function *findpeaks*, to detect maxima and minima with a prominence of 3% and a minimum peak distance of 50 frames. The rate of intensity increase per cell was measured as intensity increase over the time between the first minimum to the closest maximum. If there was only one peak detected the rate is calculated as intensity increase over the time between the peak and the local maxima or minima. If there were no peaks detected, the rate is calculated as the rate of signal increase over the minimum to the maximum of the entire trace. Every cell was therefore assigned a rate of intensity increase in relation to the maximum intensity of that cell.

