## Supplementary Figures 1-9 for "Redistribution of proteasomes through diffusion and cytoskeleton-dependent mechanisms upon stress induced by protein aggregates"

### Supplementary Figure 1

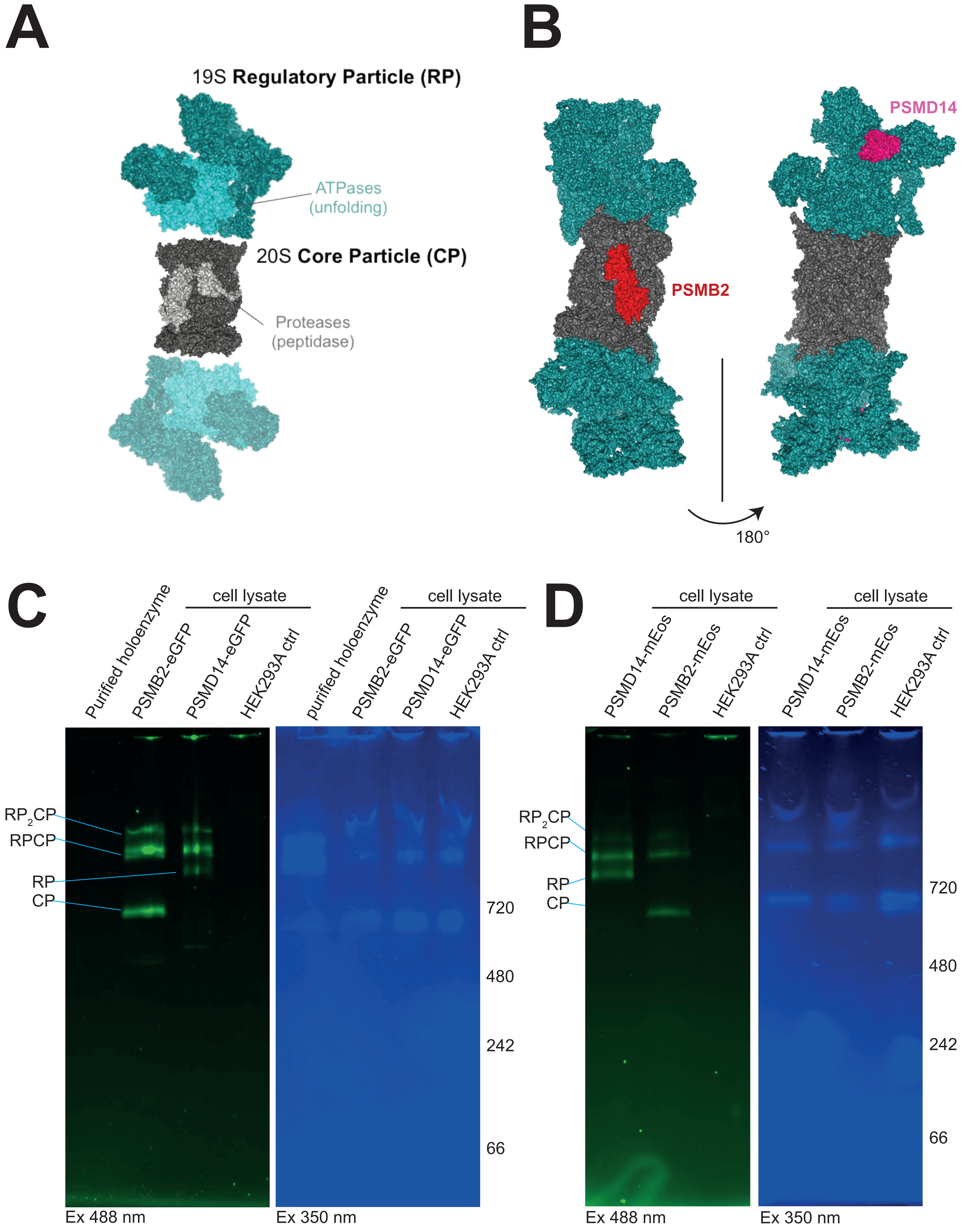

### Supplementary Figure 2

A

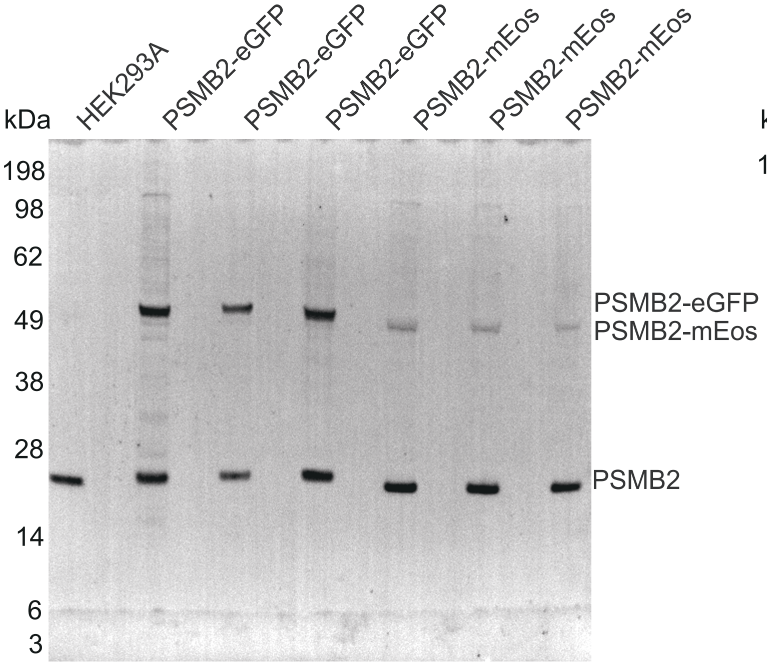


B

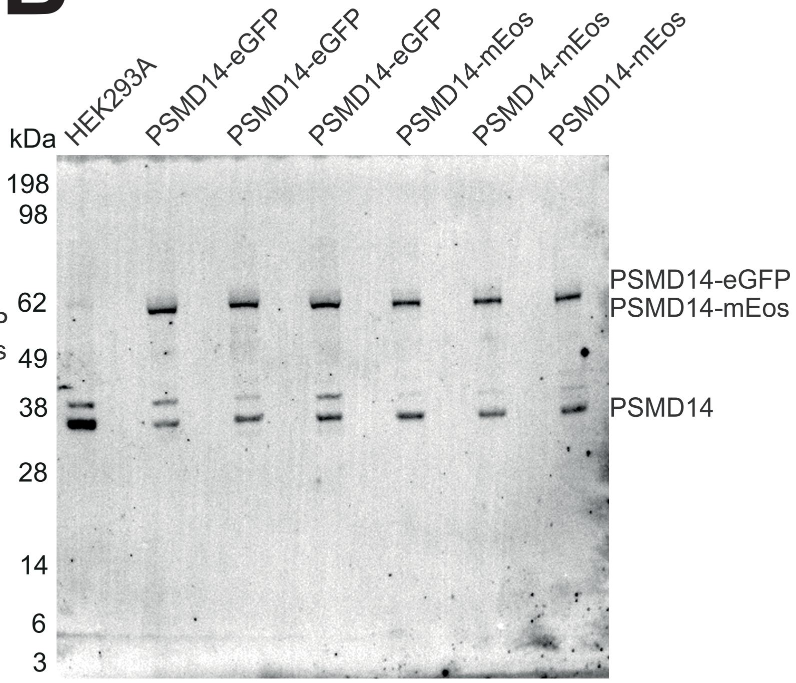


C

| Cell line | Labeling percentage |
|---|---|
| PSMB2-eGFP | 48% |
| PSMB2-mEos | 12% |
| PSMD14-eGFP | 59% |
| PSMD14-mEos | 52% |

### Supplementary Figure 3

A

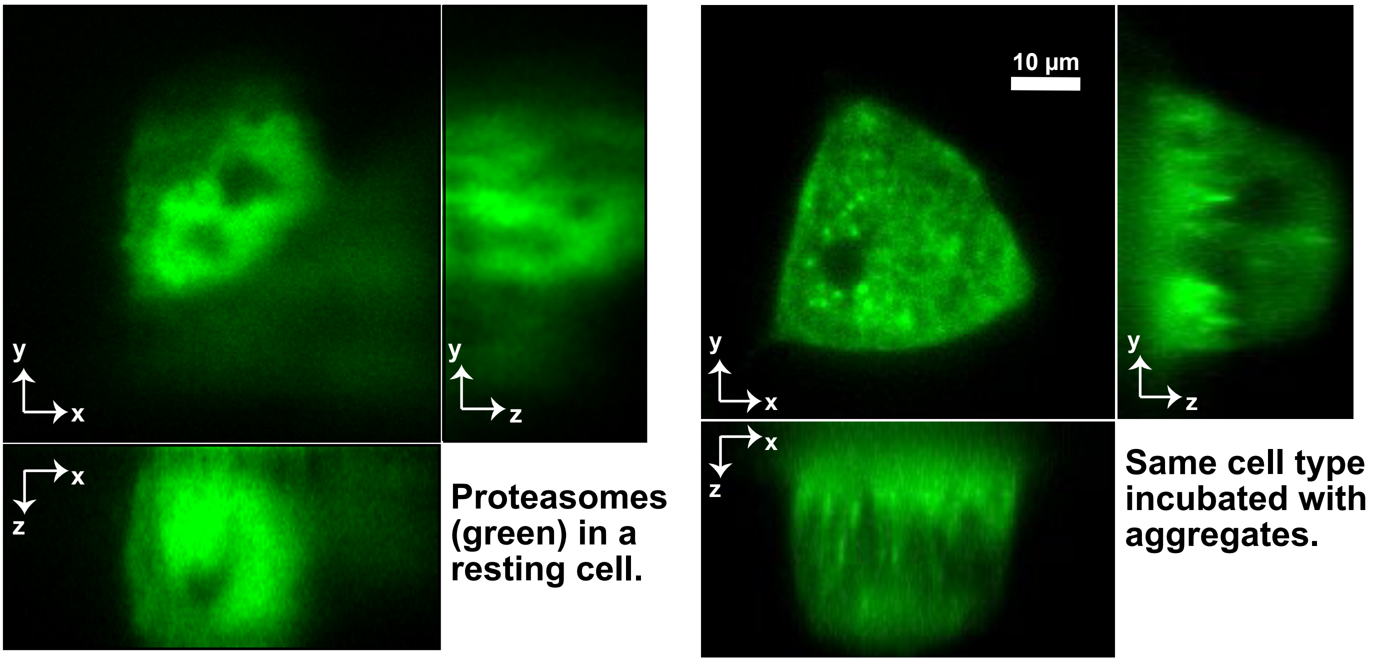


B

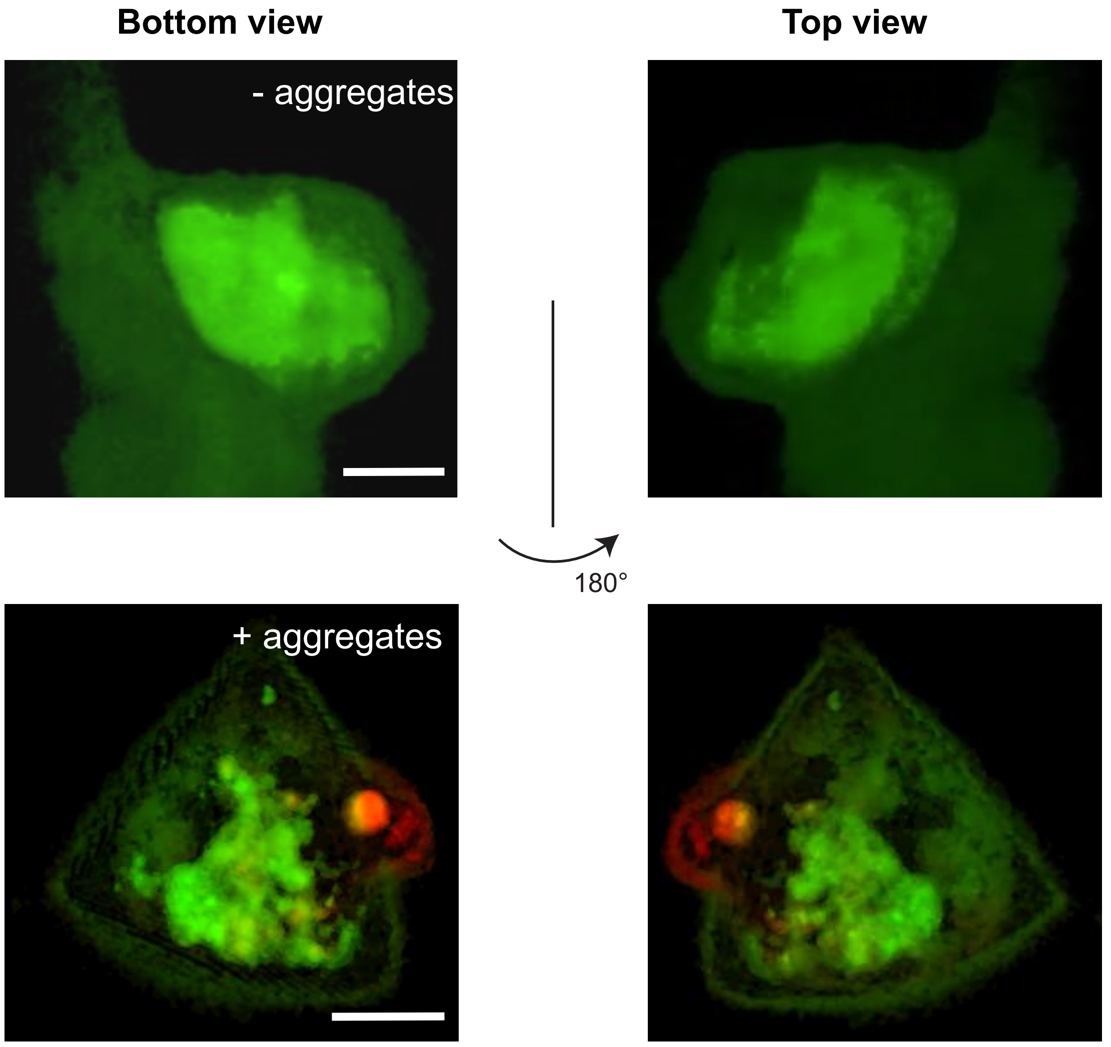

### Supplementary Figure 4

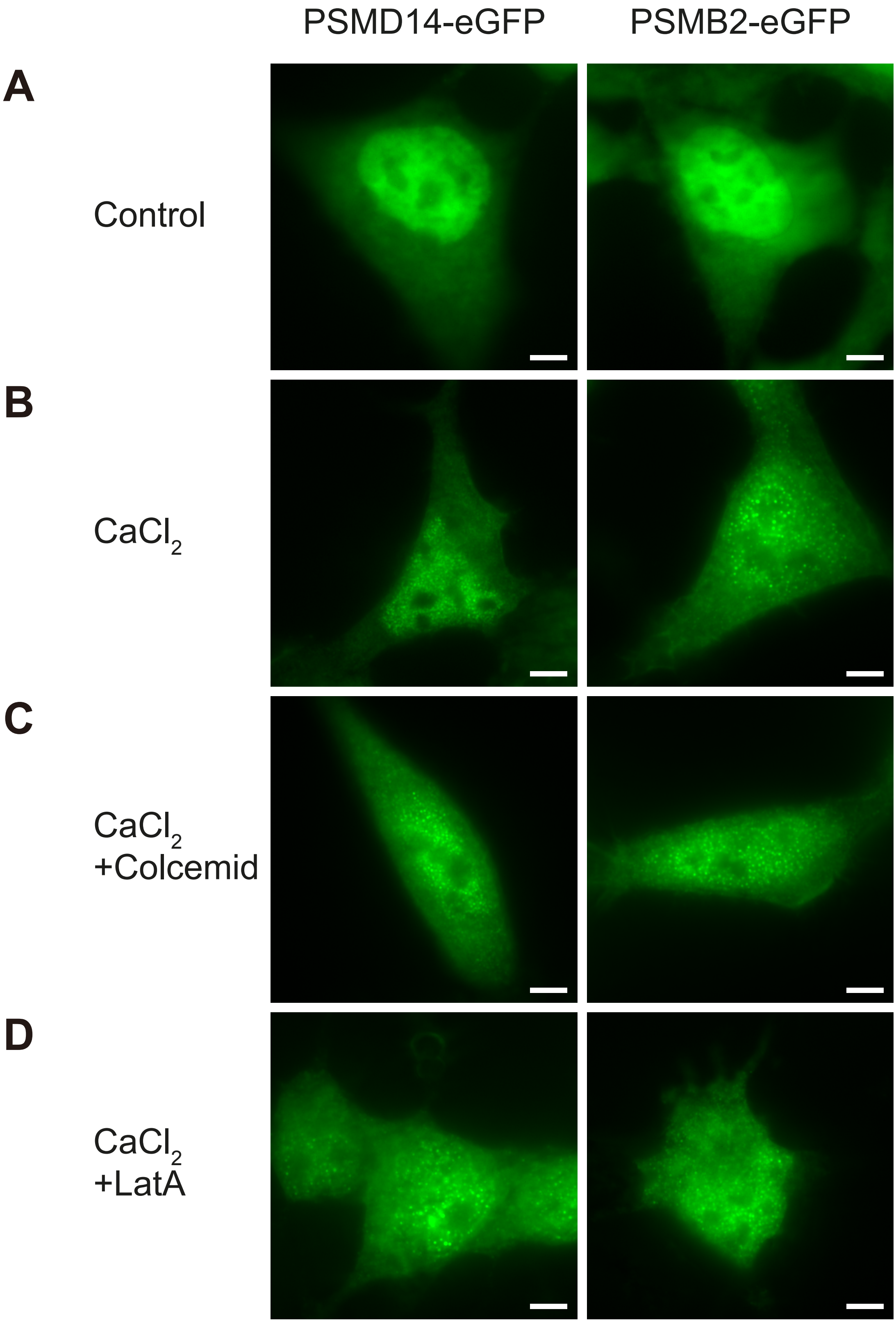

### Supplementary Figure 5

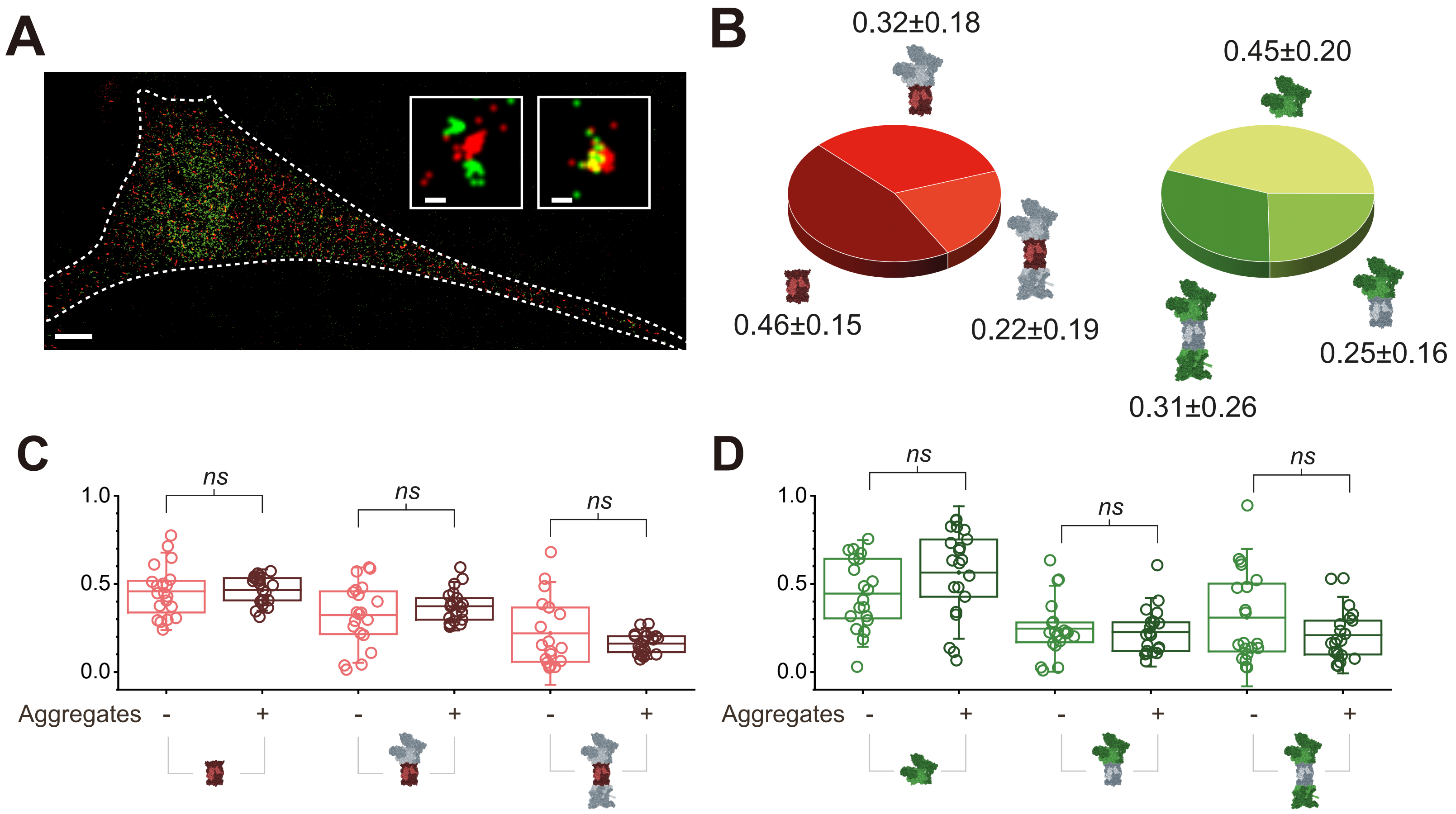

### Supplementary Figure 6

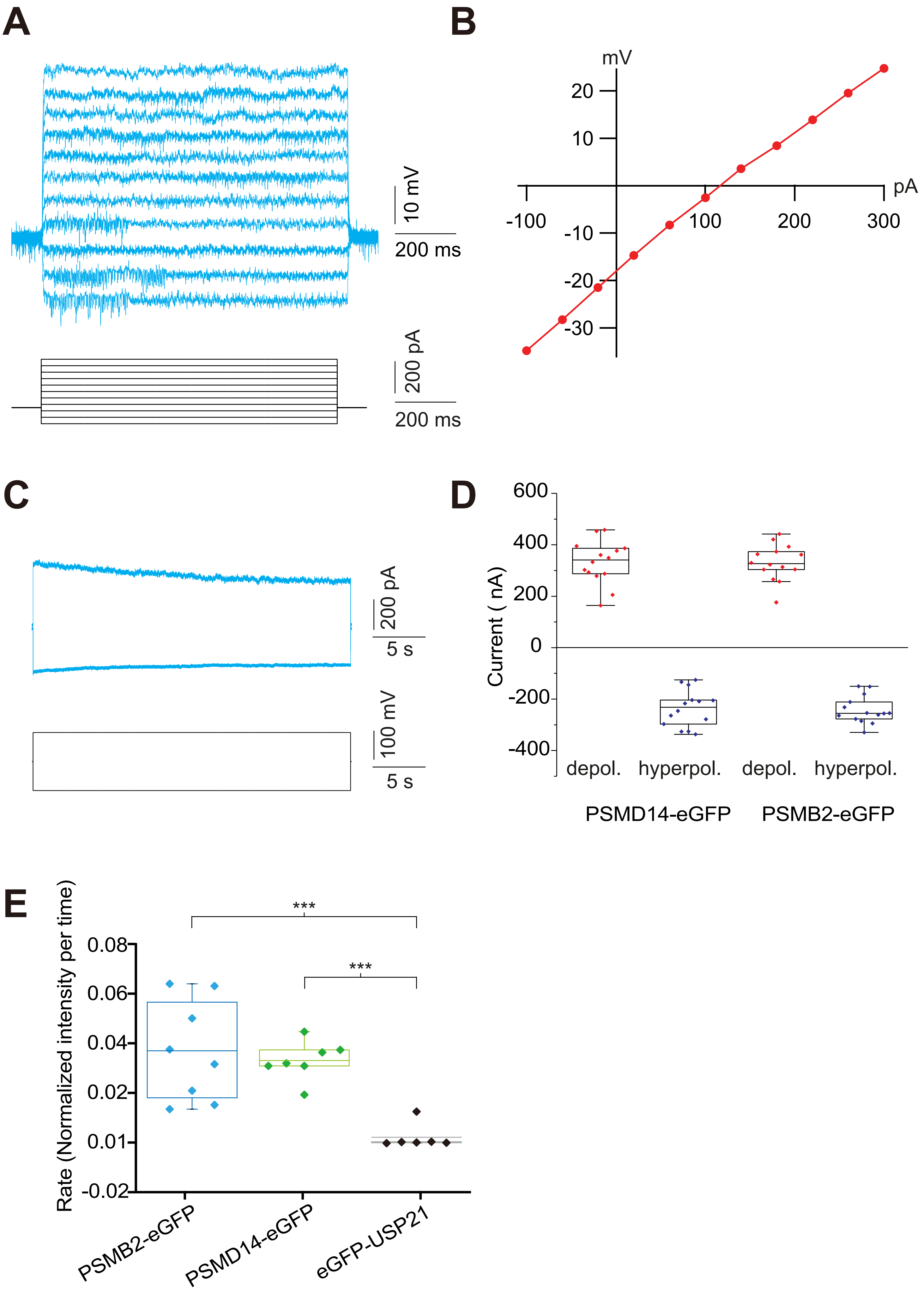

### Supplementary Figure 7

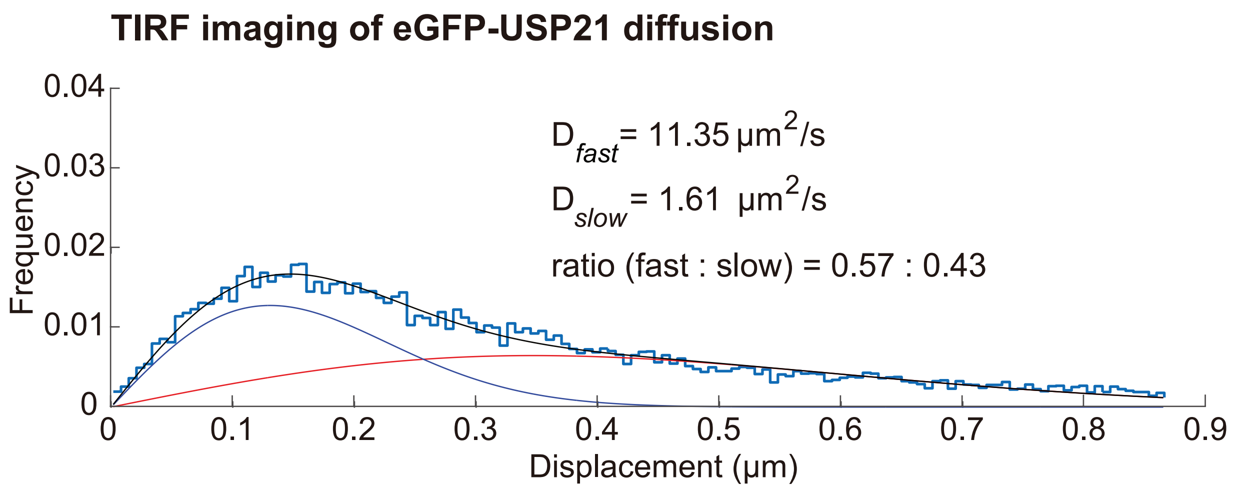


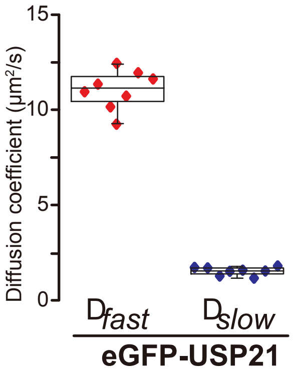


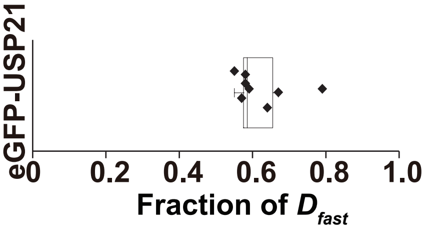

### Supplementary Figure 8

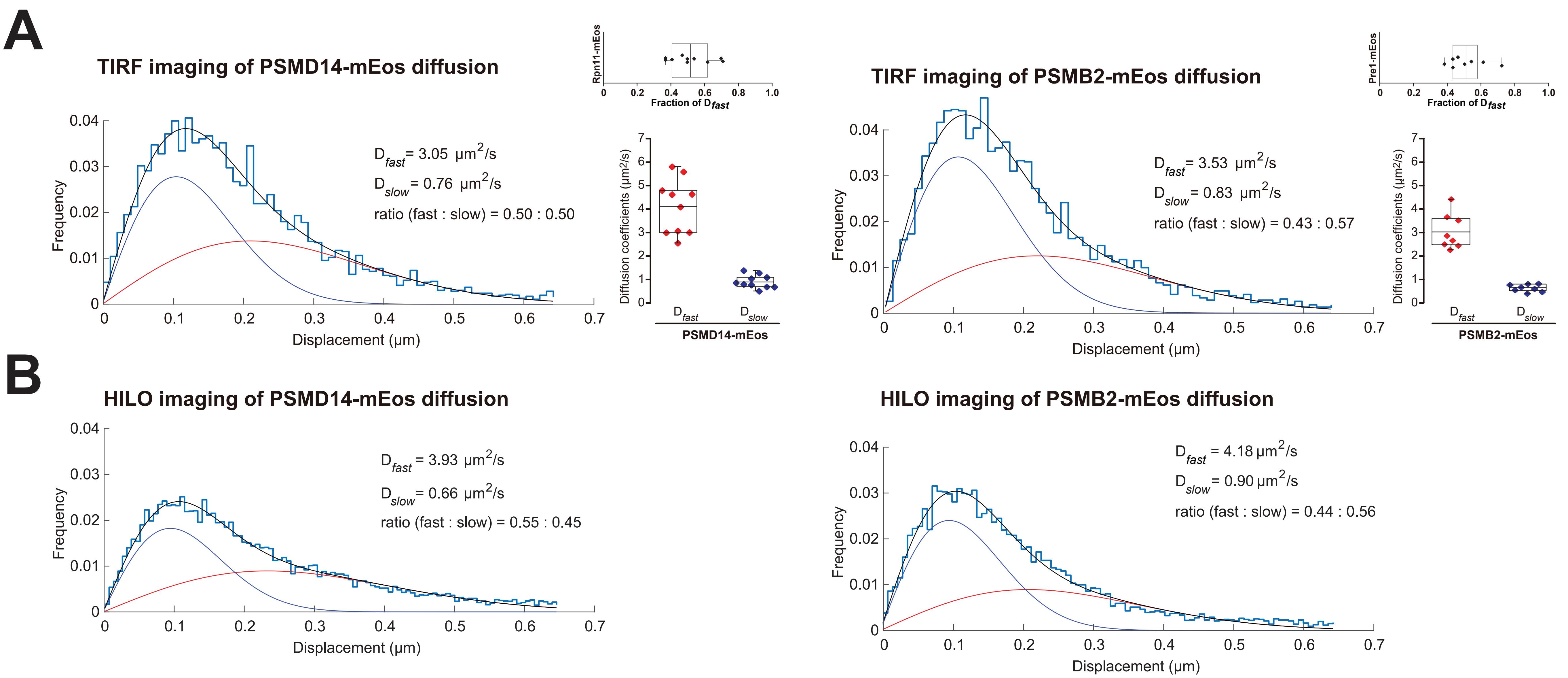

### Supplementary Figure 9

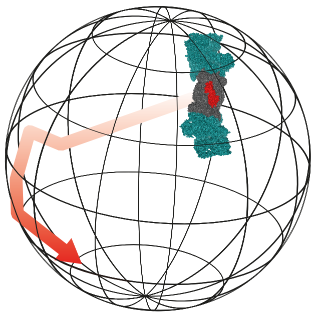

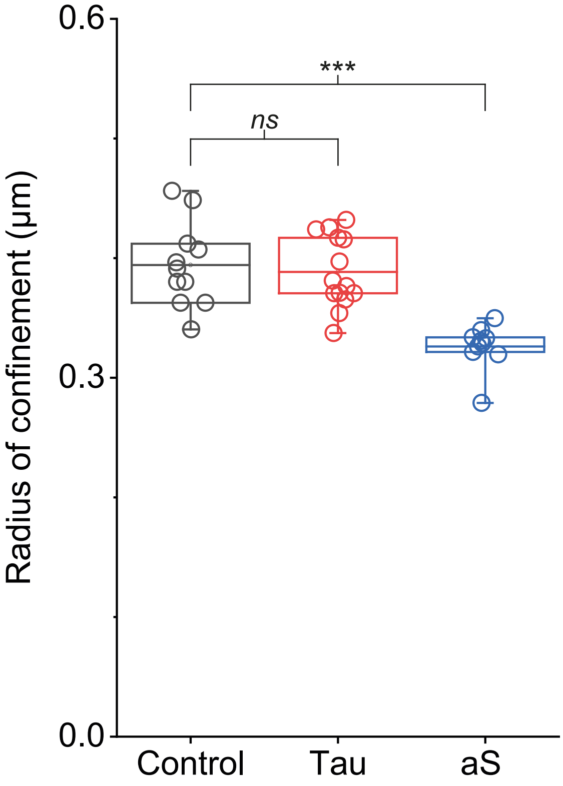
